# Genome assembly of *Spinifex sericeus* provides insights into repetitive content and haplotype variation in a dioecious coastal grass

**DOI:** 10.64898/2025.12.11.692453

**Authors:** Stephanie H Chen, Jason G Bragg, Ashley Jones

## Abstract

*Spinifex sericeus* (beach spinifex) is a perennial, dioecious grass native to coastal ecosystem of Oceania that has an important role in dune stability. We present a near telomere-to-telomere, haplotype-resolved nuclear genome alongside complete organelle genomes of *S. sericeus*. The nuclear genome is highly repeat-rich, dominated by long terminal repeat (LTR) retrotransposons. Comparative analysis of the two haplotypes reveals extensive structural variation and high levels of duplication that are suggestive of active and expanding LTRs that may drive genome evolution and expansion in this species. The chloroplast genome exhibits heteroplasmy, having two distinct circular configurations spanning 138 kbp each. The complex mitochondrial genome consists of a large linear component spanning 1.85 Mbp and a small circular 134 kbp component. These genomic resources provide a foundation for advancing research on sex determination and stress adaptation in grasses, and well as practical applications in crop improvement and genetically informed coastal dune restoration.

**Significance statement:** The reference genome of *Spinifex sericeus* provides a foundational resource for understanding adaptations to the harsh coastal dune environment, including saline conditions, extreme temperatures, and drought. The genome enables comparative studies within Poaceae, where current reference genomes are dominated by crop species, and supports conservation and restoration efforts for coastal ecosystem management.

## Introduction

Grasses (Poaceae) represent one of the largest plant families, comprising c. 11,800 species across 791 genera (Soreng et al. 2022). Poaceae dominate terrestrial ecosystems and include the world’s most important cereal crops, positioning it as both an ecologically and economically critical lineage for genomic research. Despite the challenges posed by their characteristically large and repetitive genomes, Poaceae has emerged as a model clade in plant genomics (Buell 2009). Advances in long-read sequencing technology, coupled with decreasing costs, have significantly accelerated research in Poaceae genomics, making high quality telomere-to-telomere (T2T) genomes accessible (Xie et al. 2024). However, sequencing efforts have primarily focused on agriculturally important species such as rice, maize, wheat, and sorghum. For example, rice (*Oryza sativa* L.) was the first crop sequenced (International Rice Genome Sequencing Project 2005), leading to improvements in crop breeding and production. Wild grass relatives remain a largely unexplored genomic resource, having evolved adaptive mechanisms to abiotic stresses that are increasingly valuable for crop improvement under climate change (Hultgren et al. 2025).

*Spinifex sericeus* is a dioecious perennial grass occurring on coastal foredunes in its native range of eastern Australia, New Zealand, New Caledonia, and Tonga. As a pioneer dune species, this grass survives in hostile environments (Maze & Whalley 1992) through adaptations to high salinity and heat, but faces potential threats from human disturbance, grazing (Ramsey & Engeman 1994), and diseases such as smut (Kirby 1988). *Spinifex sericeus* stabilises the frontal dune area and is valuable for dune restoration, improving both the landscape and ecological integrity in coastal regions (Jenks & Brake 2001).

Sequencing the genome of *S. sericeus* will enhance our understanding *Spinifex* biology and expand the genomic resources available for Poaceae beyond crop species. In this study, we present the first assembly and annotation of a near T2T complete haplotype-resolved nuclear genome for *S. sericeus*, and the complete chloroplast and mitochondrial genomes, using PacBio HiFi long-read sequencing data, Oxford Nanopore Technologies (ONT) ultra-long reads, and Hi-C data.

## Results and discussion

We sampled a *Spinifex sericeus* plant in its native range in Australia (Figure 1A–B). Sequencing yielded 91.7 Gbp PacBio HiFi reads, 192.1 Gbp ONT reads, and 76.2 Gbp Hi-C reads (Table S1). All HiFi reads were used as input into assembly (46x coverage), whereas ONT reads were filtered for ≥20 kbp and ≥Q10, resulting in 151.9 Gbp of input reads (76x coverage). After Hi-C read trimming and quality filtering (≥20 bp and ≥Q20), there were 74.5 Gbp of data (37x coverage). Using hifiasm (UL), the initial *de novo* assemblies had 490 contigs (haplotype 1) and 122 contigs (haplotype 2) with N50 of 195 Mbp and 186 Mbp, respectively (Table S2 and Figure S1). Hi-C scaffolding with YaHS made 13 joins, reducing the number of contigs to 485 for haplotype 1 and 111 for haplotype 2; N50 increased to 222 Mbp and 233 Mbp, respectively (Table S3). The scaffolding results also validated a hifiasm (UL) test run which incorporated Hi-C data via the --dual-scaf option.

**Figure 1.**
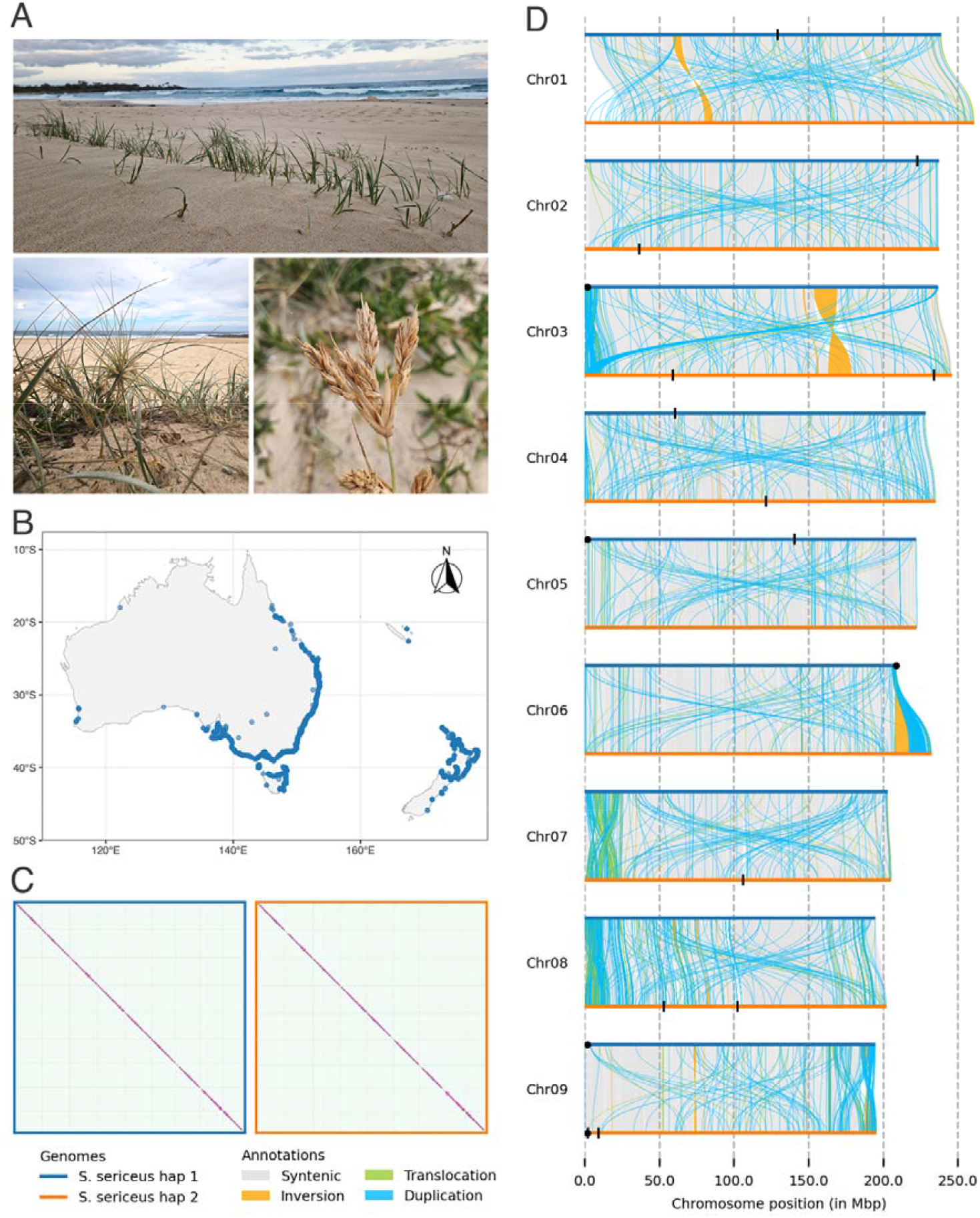
**A** *Spinifex sericeus* growing at Kioloa, New South Wales, Australia (top; photograph by Ashley Jones) and close up of flowers at Maroubra, New South Wales (female on bottom left and male on bottom right; photographs by Jason Bragg). **B** Map of Global occurrence records from the Biodiversity Information Facility (GBIF.org 2025). **C** Hi-C contact maps of haplotype 1 (left) and haplotype 2 (right) visualised in PretextView with grey lines separating scaffolds. **D** Synteny and structural variation between the two haplotypes of the *Spinifex sericeus* genome. Haplotypes 1 and 2 were aligned and genomic regions were classified as syntenic, inverted, translocated, duplicated, or haplotype specific (remaining white). Only structural variants ≥10 kbp are visualised. Vertical black lines on the genome represent gaps, where scaffolding occured (13 gaps total). Circular black markers represent missing telomeres (31/36 present).

We generated a near T2T complete haplotype-resolved reference genome for *Spinifex sericeus*, with each haplotype comprising 9 chromosome-length scaffolds (Table 1 and Figure 1C). The species has a 2C value of 5.41 pg with 2*x* = 18 (Murray et al. 2005), which approximates the haploid genome size at 2.6 Gbp. Our assembly was smaller at 1.97 Gbp for haplotype 1 and 2.04 Gbp for haplotype 2 and we confirmed diploidy via Smudgeplot (Figure S2). Both haplotypes had very high completeness scores of 99.7% and contained approximately 47,000 protein coding genes with a proteome completeness of 99.2%. Haplotype 1 contained a notably higher number of rRNAs compared to haplotype 2 (2,409 vs 86), while tRNA numbers were identical at 485 (Table 1).

**Table 1.** Summary of the *Spinifex sericeus* genome assembly, annotation, and quality assessment. Karyotype is 2n = 18.

| Statistic | Haplotype 1 | Haplotype 2 |
| --- | --- | --- |
| Total length (bp) | 1,966,204,384 | 2,039,249,938 |
| Scaffolds | 9 | 9 |
| N50 (bp) | 222,255,665 | 232,667,679 |
| Telomeres | 14/18 | 17/18 |
| T2T contigs | 2/9 | 3/9 |
| T2T scaffolds | 5/9 | 8/9 |
| Gaps | 4 | 9 |
| GC (%) | 46.35 | 46.43 |
| Genome completeness and (duplicated) | 99.7% (2.3%) | 99.7% (2.2%) |
| Protein-coding genes (GeMoMa) | 47,411 | 47,342 |
| mRNAs | 65,517 | 65,366 |
| rRNAs | 2,409 | 86 |
| tRNAs | 485 | 485 |
| Proteome completeness and (duplicated) | 99.2% (34.4%) | 99.2% (34.6%) |
| Repeats (%) | 82.06 | 82.30 |
| QV | 61.79 | 62.33 |

The *S. sericeus* genome contained a high proportion of repeats at over 82%, dominated by long terminal repeat (LTR) retrotransposons, notably Gypsy and Copia elements (Table S4); this number was higher than the estimate given by GenomeScope using both HiFi (57%) and ONT data (57%) (Figure S3). Using both Tidk (Figure S4) and seqtk (Table S5), 31 out of 36 telomeres (motif AAACCCT) were identified (summarised in Table S6). Haplotype 1 consisted of 5/9 T2T scaffolds and haplotype 2 consisted of 8/9 T2T scaffolds; there are only 13 gaps in total (Table S7). The *S. sericeus* genome had a high assembly quality value (QV) of 62 for each haplotype, indicating a base accuracy of >99.9999%, which exceeds human T2T genome projects (Cheng et al. 2025). This also highlights how long-read sequencing accuracy and *de novo* assembly for non-model species has improved substantially (Ferguson et al. 2022).

To assess the conservation of genome structure and identify structural variations, the two haplotypes were aligned and compared. Both haplotypes were highly syntenic, with 92% of haplotype 1 (1,846 Mbp) and 89% of haplotype 2 (1,847 Mbp) showing conserved sequence arrangement (Figure 1D and Table S8). The remaining sequences contained substantial structural variations distributed across the genome. Duplications were the most prominent structural variant, comprising 8,517 events in haplotype 1 totalling 58 Mbp (2.89%) and 9,318 events in haplotype 2 totalling 75 Mbp (3.60%). We observed duplications genome-wide, including inter-haplotype events spanning entire chromosome arms, potentially reflecting underlying genome organization. The majority of duplications ranged between 5 and 20 kbp, consistent with the sizes of annotated LTR retrotransposons including Gypsy and Copia elements, suggesting that transposable element activity is a key driver of genomic structural variation. This prevalence of duplications aligns with emerging evidence that they serve as a critical evolutionary mechanism in Poaceae, enabling rapid adaptation to diverse environmental landscapes (Ahrens et al. 2020; Bellec et al. 2023).

Haplotype-specific sequences represented 35 Mbp (1.72%) in haplotype 1 and 71 Mbp (3.40%) in haplotype 2, comprising novel insertions, deletions, and highly diverged regions. These diverged regions may also include duplications that have accumulated mutations beyond recognition (Ferguson et al. 2024). Translocations were moderately abundant, with 1,412 events spanning approximately 42 Mbp (2.08%) in haplotype 1 and 41 Mbp (1.97%) in haplotype 2. Inversions were rare but large, with only 95 events detected that cumulatively affected 35 Mbp (1.75%) of haplotype 1 and 46 Mbp (2.23%) of haplotype 2. Read mapping to both haplotypes and subsequent coverage analysis verified the structural variants, haplotype-specific regions, and confirmed high confidence in the genome assembly quality (Figure S5). The extensive structural variation between haplotypes underscores the remarkable genome evolution within this species and suggests that structural variants contribute to genomic diversity and adaptive potential.

We also successfully assembled complete chloroplast and mitochondrial genomes, revealing complex structural arrangements in both organelles. The *S. sericeus* chloroplast exhibited heteroplasmy, having two distinct circular configurations, of 137,659 bp each, which is consistent with other Poales grasses (Zhou et al. 2025). The mitochondrial genome consisted of two components: a considerably large linear sequence of 1,851,594 bp, and a circular sequence of 134,119 bp (Figure S6). It is possible the linear sequence may be incomplete; however, mitochondrial genomes consisting of a linear and circular component have been reported, for example *Alnus glutinosa* (Zhou et al. 2025). Similarily, Poales mitochondria genomes have been oberved to be larger on average, for example, the 2.73 Mbp mitochondria of *Carex laevigata* (Zhou et al. 2025).

This reference genome of *S. sericeus* represents a foundational resource for understanding grass genome evolution and adaptive capacity under abiotic stress. Thriving under the high salinity, temperature extremes, and intense light of coastal dunes, *S. sericeus* offers opportunities to identify genetic mechanisms underlying stress tolerance, which are critical for developing climate-resilient crops (Melino & Tester 2023). As a dioecious species, this genome also enables investigation of sex-biased gene expression across reproductive and vegetative tissues, such as inflorescences, leaves, and roots (Sanderson et al. 2019). Field studies have observed a male bias in the observed ramet sex ratio (Maze & Whalley 1990), raising questions about the underlying mechanisms driving this imbalance. Genes involved in sex determination and ecological adaptations could be investigated and contribute to genetically informed restoration strategies to enhance coastal dune resilience and conservation.

## Materials and methods

### Sampling, DNA extraction, and sequencing

Cuttings were sampled from a single plant in Yarra Bay Beach, New South Wales, Australia (33.9774° S, 151.2280° E; BioSample SAMN41648390) and propagated in a growth chamber in the original sand substrate supplemented with a seaweed solution (Seasol) fortnightly. The sex of the reference genome is currently unknown as the plant was not flowering at the time of collection, nor flowered in the 1.5 years that it has been in cultivation. Young leaf blades were harvested and cryogenically stored at -80°C for DNA extractions and Hi-C.

High-molecular weight DNA was extracted using a magnetic bead-based protocol as previously described (Jones et al. 2021). The DNA quality was assessed on a Femto Pulse system (Figure S7) using a genomic DNA 165 kbp kit (Agilent Technologies), and subsequently size selected for fragments ≥20 kbp using a BluePippin (Sage Science). A Pacific Biosciences (PacBio) DNA library was prepared using the SMRTbell prep kit 3.0 and sequenced on the Revio platform using a 25M SMRT cell, to generate highly accurate HiFi reads (≥Q20). Two Oxford Nanopore Technologies (ONT) native DNA libraries were prepared using the Ligation Sequencing Kit V14 (SQK-LSK114) and sequenced on the PromethION P24 platform using two FLO-PRO114M R10.4.1 flow cells, to generate ultra-long reads. The ONT flow cells were washed, re-primed, and reloaded twice when sequencing declined (ONT Flow Cell Wash Kit EXP-WSH004). ONT sequencing data (pod5) was basecalled to fastq with ONT Dorado v7.4.12 using the super-accurate DNA model (sup v4.3.0, e8.2, 400 bps). The ONT fastq reads were filtered for ≥20 kbp and ≥Q10 using chopper v0.8.0 (De Coster et al. 2018).

A chromosome conformation Hi-C library was prepared using a Phase Genomics Proximo Hi-C Plant Kit v4 (KT3040B) and sequenced on an Illumina NovaSeq 6000 using an S4 flow cell with a 300 cycle kit (150 bp paired-end sequencing). Hi-C reads were trimmed for Illumina adapters (--nextera) and filtered for ≥20 bp and ≥Q20. Read pairs were validated with Trim Galore v0.6.2 (Krueger et al. 2023).

### Genome assembly

Independently using PacBio HiFi reads and filtered ONT ultra-long reads, the nuclear genome was profiled by k-mer frequencies (*k* = 31) to assess genome size, heterozygosity and ploidy, with FastK v1.1, GenomeScope v2.0.1, and Smudgeplot v0.4.0 (Ranallo-Benavidez et al. 2020). A *de novo* haplotype-resolved genome assembly was performed with hifiasm (UL) v0.24.0-r703 (Cheng et al. 2024), using PacBio HiFi reads, filtered ONT ultra-long reads (--ul), filtered Illumina Hi-C reads (--h1 --h2), and specifying the representative Poaceae telomere motif (--telo-m AAACCCT). To scaffold the genome, Hi-C reads were mapped with HiC-Pro v3.1.0 (Servant et al. 2015) and scaffolding was performed with YaHS v1.2.2 (Zhou et al. 2023). Hi-C contact maps were generated using PretextMap v0.1.9 (Harry 2022) and visualised with PretextView v1.0.5 (Harry & Guan 2025). Gaps of unknown length were standardised to 100 bp. Scaffolds less than 1 Mbp were filtered out so that the remaining sequences were the nine nuclear chromosomes. The nine chromosomes were ordered by length, according to haplotype 1.

To assemble organelle genomes, the PacBio HiFi reads were first randomly subsampled to 2 M reads using seqtk sample v1.5 (r133) (Li 2025). Approximately 5% of reads were organelle, capturing thousands of times coverage of each organelle genome. The chloroplast genome was assembled with TIPPo v1.3.0 (Xian et al. 2025) and mitochondria genome assembled with OatK v1.0 (Zhou et al. 2025).

### Assembly quality assessment

All genome assembly graphs were inspected with Bandage v0.8.1 (Wick et al. 2015). Nuclear genome completeness was assessed by Benchmarking Universal Single-Copy Orthologs (BUSCO) with the Poales OrthoDB v12 database (*n* = 6,282), using compleasm v0.2.7 (Huang & Li 2023). To assess base accuracy of the genome, yak v0.1 (r56) was used to compare k-mer spectra (*k* = 31) between the assembly and PacBio HiFi reads to calculate a coverage-adjusted assembly quality value (QV) (Cheng et al. 2021). To assess genome coverage, PacBio HiFi reads and filtered ONT ultra-long reads were independently mapped to the genome using Minimap2 v2.29 (Li 2018) and coverage was plotted using the JVarkit v2024.08.25 tool WGSCoveragePlotter (Lindenbaum 2024).

### Genome annotation

Genes were annotated with GeMoMa v1.9 (Keilwagen et al. 2019) with five RefSeq genomes: rice (*Oryza sativa*; GCF_034140825.1) (Shang et al. 2023), corn (*Zea maize*; GCF_902167145.1) (Hufford et al. 2021), common reed (*Phragmites australis*; GCF_958298935.1) (Christenhusz et al. 2024), green foxtail (*Setaria viridis*; GCF_005286985.2) (Thielen et al. 2020), and *Arabidopsis thaliana* (TAIR 10.1; GCF_000001735.4) (Lamesch et al. 2012). Completeness was assessed with compleasm v0.2.7 (Huang & Li 2023) in protein mode with the Poales OrthoDB v12 database. The rRNA genes were predicted with Barrnap v0.9 (Seemann 2018) and tRNAs were predicted with tRNAscan-SE v2.0.12 (Lowe & Chan 2016), applying the high confidence filter.

Repeats were annotated with a custom library using RepeatModeler v2.0.2a with - engine ncbi (Flynn et al. 2020) and RepeatMasker v4.1.2 (Tarailo□Graovac & Chen 2009). Telomeric repeats were identified with tidk v0.2.65 (Brown et al. 2025) and seqtk v1.5 (r133) (Li 2025).

### Synteny and structural variation between haplotypes

To assess nuclear genomic differences, the two haplotypes were aligned to each other with the MUMmer4 v4.0.1 tool nucmer (--maxmatch -l 50 -b 500 -c 200) (Marçais et al. 2018). All shared 50-mers between haplotypes were identified and adjacent 50-mers were merged into a single alignment. Alignments were filtered for

≥200 bp and ≥95% sequence identity with MUMmer4 tool delta-filter. Synteny and genomic rearrangements were identified with SyRI v1.6.3 (Goel et al. 2019), and results ≥10 kbp were visualised with plotsr v1.1.1 (Goel & Schneeberger 2022).

## Author contributions

**Stephanie Chen** – Formal Analysis, Investigation, Methodology, Software, Writing – Original Draft Preparation, Writing – Review & Editing. **Jason Bragg** – Investigation, Writing – Review & Editing. **Ashley Jones** – Conceptualization, Formal Analysis, Investigation, Methodology, Software, Visualization, Writing – Original Draft Preparation, Writing – Review & Editing.

## Data accessibility statement

The *Spinifex sericeus* reference genome and raw sequencing data (PacBio HiFi, ONT, and Hi-C data) were deposited to NCBI under BioProject PRJNA1119262 and PRJNA1346877 embargoed until acceptance of paper. Additional data supporting this work, including genome annotation files, are available on the CSIRO Data Access Portal (https://doi.org/10.25919/2esb-qs53).

## Acknowledgements

We acknowledge Australia’s First Nations Peoples as the Traditional Custodians of the land and waters on which this work was conducted, and pay our respects to all Elders past, present, and emerging. We would like to thank the Biomolecular Resource Facility at the John Curtin School of Medical Research, ANU in Canberra, Australia, where sequencing was conducted. This research acknowledges the support provided by NCRIS enabled Bioplatforms Australia infrastructure. This research was undertaken with the assistance of resources from the National Computational Infrastructure (NCI Australia), an NCRIS enabled capability supported by the Australian Government.

## Benefit sharing statement

All raw sequencing data and assembled genomes have been shared with the broader public via appropriate biological databases.

